# Hyaluronan Synthesis Inhibition Impairs Antigen Presentation and Delays Transplantation Rejection

**DOI:** 10.1101/2020.08.28.272443

**Authors:** Payton L. Marshall, Nadine Nagy, Gernot Kaber, Graham L. Barlow, Amrit Ramesh, Bryan J. Xie, Miles H. Linde, Naomi L. Haddock, Colin A. Lester, Quynh-Lam Tran, Christiaan de Vries, Andrey Malkovskiy, Irina Gurevich, Hunter A. Martinez, Hedwich F. Kuipers, Koshika Yadava, Xiangyue Zhang, Stephen P. Evanko, John A. Gebe, Xi Wang, Robert B. Vernon, Aviv Hargil, Carol de la Motte, Thomas N. Wight, Edgar G. Engleman, Sheri M. Krams, Everett Meyer, Paul L. Bollyky

## Abstract

A coat of pericellular hyaluronan surrounds mature dendritic cells (DC) and contributes to cell-cell interactions. We asked whether 4-methylumbelliferone (4MU), an oral inhibitor of HA synthesis, could inhibit antigen presentation. We find that 4MU treatment reduces pericellular hyaluronan, destabilizes interactions between DC and T-cells, and prevents T-cell proliferation *in vitro* and *in vivo*. These effects were observed only when 4MU was added prior to initial antigen presentation but not later, consistent with 4MU-mediated inhibition of *de novo* antigenic responses. Building on these findings, we find that 4MU delays rejection of allogeneic pancreatic islet transplant and allogeneic cardiac transplants in mice and suppresses allogeneic T-cell activation in human mixed lymphocyte reactions. We conclude that 4MU, an approved drug, may have benefit as an adjunctive agent to delay transplantation rejection.

## Introduction

The pericellular matrix (PCM) or glycocalyx is a “coat” of proteoglycans and protein-associated polysaccharides that extends outwards from the cell membrane(1–4). While the PCM is best characterized in regards to chondrocytes(5) and endothelial cells(6), PCM-like structures surround many cell types(7–9) and contribute to osmotic balance(10), migration(11), and diffusion of nutrients and signaling molecules(12). The PCM is an important element of cellular function and a mechanism to interact with the environment beyond the cell surface.

In chondrocytes and endothelial cells, the PCM contributes to cell-cell communication(13– 15). These interactions are mediated via dynamic, non-covalent molecular interactions between matrix components and receptors(16–19) on adjacent cells. In addition, interactions between opposing pericellular matrices themselves may generate adhesive forces akin to the self-adhesion of dynamically cross-linked hydrogels(20).

The PCM surrounding leukocytes consists mainly of hyaluronan (HA)(1, 21, 22), a repeating disaccharide of N-acetylglucosamine and D-glucuronic acid(23). HA is synthesized by three HA synthase (HAS) isoforms, HAS1, HAS2 and HAS3(24), and is extruded directly through the plasma membrane to the extracellular space(25). Additionally, a number of proteins and proteoglycans crosslink HA on the cell-surface and create higher order levels of structure that may be significant to the stability and function of the PCM coat(26, 27).

Pericellular HA may contribute to dendritic cell (DC)/T-cell interactions. We previously reported that HA is present in focal deposits in association with MHC-II at the immune synapse (IS), the interface between DC and T-cells(28). In another study, physical blocking of HA via peptide limited DC/T-cell interactions(29). HA is abundant in lymphatics(30) and deposits are present within secondary lymphatic tissues of autoimmune diabetes patients(31). However, the particular cell types that produce HA within lymph nodes (LN) and the functional contributions of HA to *in vivo* antigenic responses are unclear.

Treatment with 4-methylumbelliferone (4MU), an inhibitor of HA synthesis(32, 33), is beneficial in several models of inflammation, innate immunity, and fibrosis(34, 35). We and others have demonstrated that 4MU can prevent or treat multiple mouse models of autoimmunity(36), including diabetes(37), multiple sclerosis(38, 39), and rheumatoid arthritis(40). We previously reported that these effects were associated with enhanced Foxp3+ regulatory T-cell (Treg) homeostasis(37, 38, 41–44). As peripheral Treg induction is known to be associated with weak antigenic signals(45–47), this and the aforementioned data on pericellular HA in cell-cell interactions raised the intriguing possibility that 4MU treatment might impact antigen presentation.

In the present study we asked whether pericellular HA contributes to antigen presentation *in vitro* and *in vivo*, and whether 4MU can be used to prevent antigenic responses involved in transplant rejection.

## Results

### HA is present at sites of antigen presentation and is produced primarily by antigen presenting cells (APC)

We first sought to determine if HA is present *in vivo* at sites of antigen presentation. To investigate this, we used a well-established model of pancreatic islet inflammation – the DORmO mouse(48). To this end, we stained sections of spleen and pancreatic lymph nodes (LN) isolated from healthy adult mice (**Fig. 1A, B**) and mice with active autoimmune inflammation (**Fig. 1C, D**) for HA.

**Figure 1:**
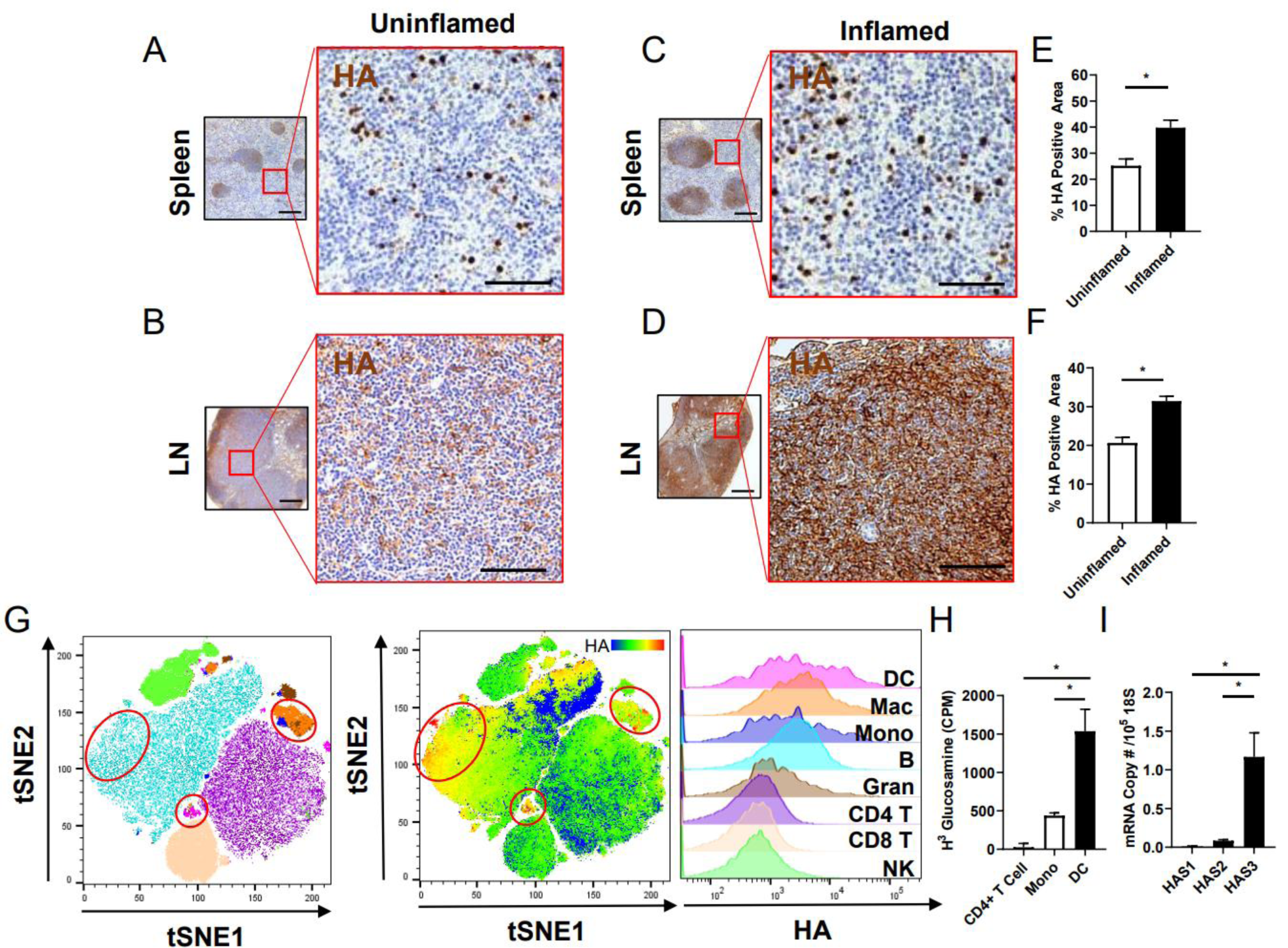
HA is produced by antigen presenting cells *in vivo*. A-F. Staining of HA in spleen (A,C) and pancreatic lymph node (LN)(B,D) sections isolated from (A,B) uninflamed DO11.10 control mice and (C,D) age/gender-matched actively inflamed mice (pre-diabetic DORmO mice). Red boxes highlight interfollicular regions, sites of antigen presentation to CD4+ T-cells. HA positive area in stimulated and unstimulated mice are shown for spleen (E) and pancreatic lymph node (PLN) (F). Data include information from >10 sections per mouse and n=3 mice per condition. G. tSNE plots representing different immune subsets present in murine lymphatic tissue including Dendritic Cells (DC)(magenta), Macrophages (Mac)(orange), Monocytes (Mono)(dark blue), B-cells (B)(light blue), Granulocytes (Gran)(brown), CD4+ T-cells (CD4 T)(purple), CD8+ T-cells (CD8 T)(pink) and NK cells (NK)(green) alongside a heat map showing the relative binding of HABP. Areas circled in red mark cell types associated with substantial cell surface HA. Histogram plots for HABP binding intensity are based on cell definitions provided in Supplementary Figure 2A and match those colors in the tSNE plot. Data are representative of n=3 separate experiments. H. Counts per minute (CPM) of radiolabeled glucosamine incorporation into HA by purified human CD4+T cells, monocytes, and mDC. CPM shown is for the amount of radiolabel lost upon treatment of cell lysates with hyaluronidase to reflect quantify only cell-bound HA. I. mRNA expression by human mDC of the three HA synthases, HAS1-3, all normalized to 18S. Data include pooled isolated from 4 different donors and measured in duplicate. Data represent mean ± SEM; *, p < 0.05 vs. respective control by unpaired t-test.

We observed that HA deposition was more pronounced in both the spleens (**Fig. 1E**) and LNs (**Fig. 1F**) of mice with active inflammation. HA deposition was most pronounced within germinal centers and interfollicular regions, typical sites of antigen presentation to B and T lymphocytes, respectively. Similar robust HA staining was seen in inflamed human tonsil sections collected from de-identified human donors (**Supplemental Fig. 1**).

To characterize the cell types associated with this increased HA expression, we analyzed disaggregated DORmO lymphatic tissues via flow cytometry. The gating schemes for our cell definitions are shown in **Supplemental Figure 2A**. Cell surface HA was quantified by binding of fluorescently labeled HA binding protein (HABP). We observed that the highest HA expression levels were in association with DCs (CD11c+IA/IE ^high^), macrophages (CD11b+F4/80+), and monocytes (CD11b+CD14-). B cells (CD19+TCRVβ-) showed moderate surface HA expression while T-cells (TCR Vβ+CD19-) and NK cells (CD49b+) showed minimal HA expression (**Fig. 1G**).

**Figure 2.**
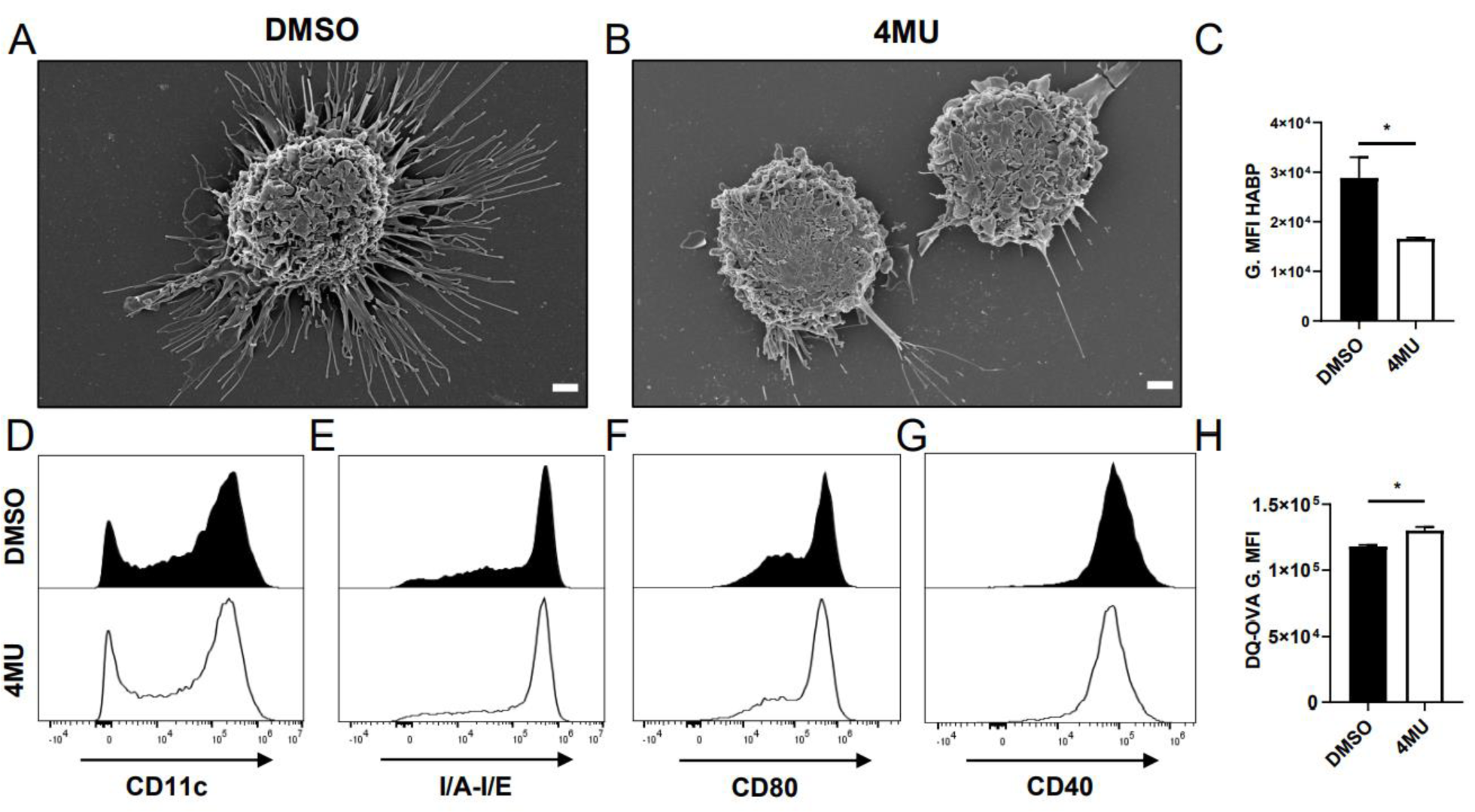
HA production alters DC phenotypes. C57Bl/6J Bone marrow derived dendritic cells (BMDC) were cultured in the presence of LPS (A) with or (B) without 4MU and the appearance of these was assessed using scanning electron microscopy (SEM). Data are representative of 2 independent experiments and >50 cells. C. Cell surface HA staining for the same cell types and conditions. D-G. Expression of cell surface markers by LPS activated murine DC in the absence or presence of 4MU, shown for (D) CD11c, and expression of (E) I/A-I/E, (F) CD80, and (G) CD40 on CD11c+ cells. H. DQ-OVA fluorescence following 30 minutes of processing after pretreatment with or without 4MU. Data for (D-M) include pooled isolated from 4 mice and measured in triplicate. Data represent mean ± SEM; *, p < 0.05 vs. respective control by one-way ANOVA with Bonferroni multiple comparisons.

To evaluate this biochemically and confirm this relationship in human cells, we radiolabeled the HA precursor glucosamine and allowed its incorporation into surface bound HA by isolated cell populations. The proportion of pericellular HA was calculated from parallel aliquots digested with hyaluronidase. We found that monocyte derived DC (mDC) produce substantially more HA than unstimulated monocytes (Mono) or CD4+ T-cells (**Fig. 1H**).

We next performed real time PCR on RNA transcripts isolated from these same cells to identify which HAS isoform predominantly is expressed during inflammation. We found that HAS3 is the most highly expressed HAS isoform in DCs (**Fig. 1I**). This is consistent with our previous finding that HAS3 is the most abundant HAS transcript in circulating mouse leukocytes(28).

Together, these data suggest that HA is present *in vivo* at sites of antigen presentation in both human and murine tissues and that antigen presenting cells are among the highest expressers of surface bound HA.

### 4MU treatment alters DC morphology but not costimulatory molecules or antigen processing

We next asked how surface HA contributes to DC phenotypes. We first performed scanning electron microscopy on DC treated with DMSO vehicle control (**Fig. 2A**) or 4MU (**Fig 2B**). HA inhibition strongly reduced spreading and the formation of dendrites - phenotypes associated with DC antigen uptake and presentation(49).

We next examined cell-surface HA expression on DC using flow cytometry (a gating scheme is shown in **Supplemental Figure 2B)**. 4MU treatment substantially decreased cell surface HA as assessed by binding of fluorescently labeled HABP (**Fig. 2C**), but showed negligible changes in CD11c, I/A-I/E, CD80, and CD40 expression relative to DMSO-treated DC (**Fig 2D-G**). These data suggest that inhibition of HA synthesis with 4MU does not adversely impact MHC-II expression or maturation as measured by costimulatory surface markers.

To assess the impact of 4MU on antigen processing, we examined the ability of DC to take up and digest DQ-OVA, a molecule that becomes fluorescent within the lower pH of endosomal vesicles. 4MU did not decrease DQ-OVA processing compared to DMSO control and indeed showed a modest increase (**Fig. 2H)**. These data indicate that 4MU does not adversely affect antigen uptake and processing.

### HA contributes to interactions between T-cells and DC

We next asked whether HA contributes to interactions between DC and T-cells. To assess this, we cultured DC and CD4+ T-cells together with polyclonal stimulus (anti-cd3/cd28 antibodies) in the absence or presence of 4MU (**Fig. 3A, B**). This allows DC to stimulate CD4+ T-cells in an antigen nonspecific manner via direct presentation of the antibodies by the DC to T-cell. We observed that HA surrounds clusters of activated, proliferating cells and that focal deposits of HA colocalize with areas dense with MHC-II (**Fig. 3C**), as we reported previously(28). 4MU decreases this HA and abrogates cell-cell interactions (**Fig. 3B, D**). To quantify the impact of 4MU on cell-cell interactions, we labeled BMDC and CD4+T-cells with SNARF-1 and CFSE, respectively and evaluated their binding interactions. We observed that 4MU decreased the number of T-cells bound to each DC (**Fig. 3E-G**) as well as the overall fraction of DCs bound to T-cells (**Fig. 3H**). These data are consistent with similar effects previously reported for PEG-Pep1 and hyaluronidase using a similar assay(29, 50).

**Figure 3:**
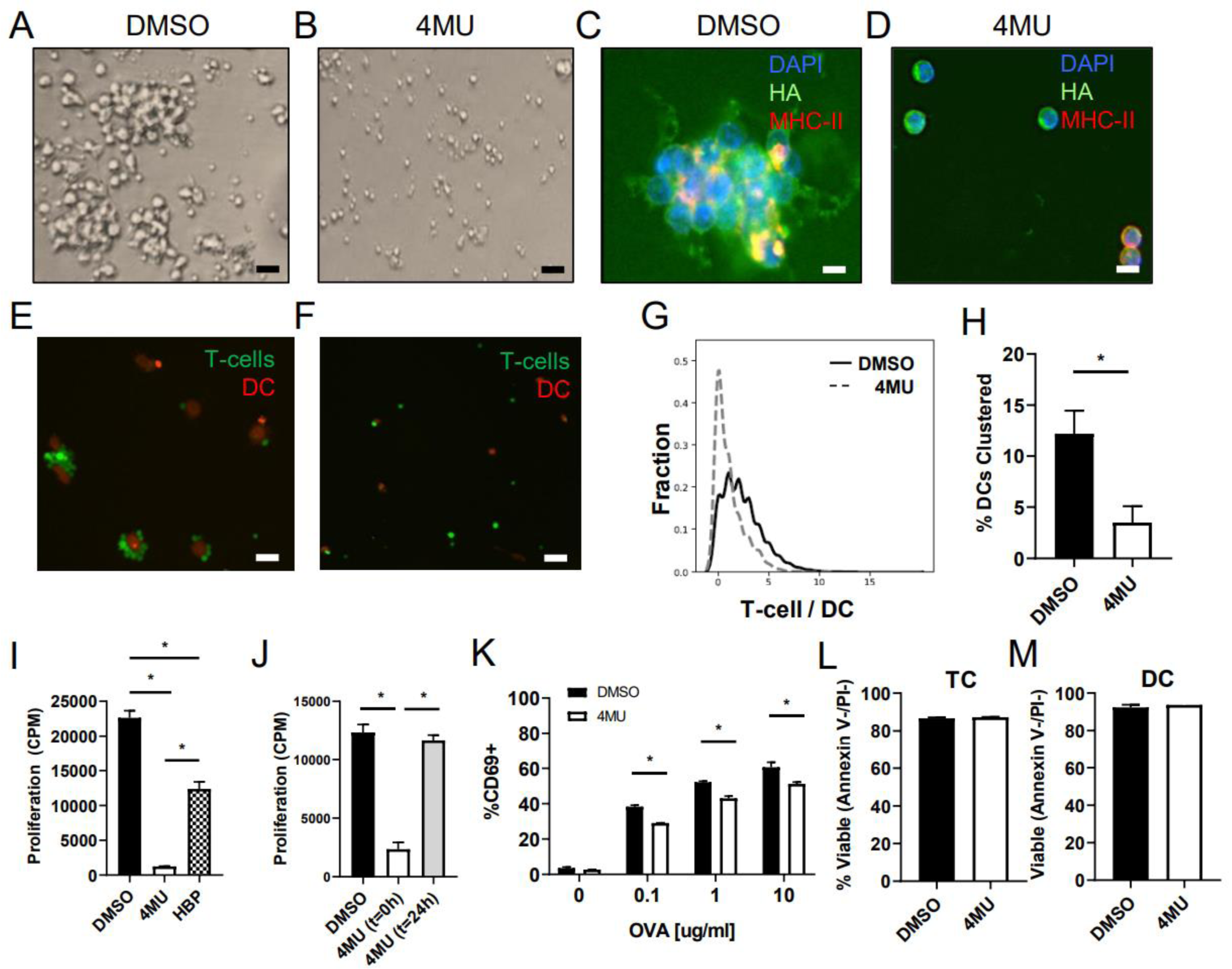
HA contributes to immune synapse formation and antigen presentation. CD4+ T-cells cultured with autologous DC pre-loaded with αCD3 and αCD28 antibodies. A-B. Appearance of cell cultures treated with (B) or without (A) 4MU. C-D. Representative images of a cultured cells treated (D) with or (C) without 4MU and stained for DAPI (blue), HA (green) and MHC-II (red). E, F. Images of DC (red) and T-cells (green) (F) with or (E) without 4MU. G-H. Quantification of the number of T-cells bound to each DC (G) and the percentage of DC engaged in binding interactions with CD4+ T-cells (H). Data are representative of two separate experiments measured in triplicate. I. Proliferation of T-cells co-cultured with APC and αCD3/αCD28 antibodies. Cells were stimulated for 72 hours with tritiated thymidine (TT) added for the final 24 hours. CPM data are for triplicate wells. J. T-cells were activated as in (I) in the setting of 4MU added at either the inception of the assay (0 hours) or 1 day later (24 hours). For the final 24 hours tritiated thymidine was added. K. %CD69+ cells of total OT-II CD4+ T-cells following activation in the setting of OVA-loaded autologous DC for five hours, showing pooled replicates for increasing concentrations of OVA. L,M. Viability of CD4+ T-cells (L) and DC (M) as assessed by annexin V and PI staining. Data for panels I-L include pooled isolated from 4 mice and are each representative of >2 experiments. Data represent mean ± SEM; *, p <0.05 vs. respective control by unpaired t-test or one-way ANOVA with multiple comparisons where appropriate.

Consistent with these effects on immune synapse formation, proliferative responses by T-cells were lost upon addition of 4MU (**Fig. 3I**). HABP likewise diminished these responses as reported previously(29) but not as effectively as 4MU (**Fig. 3I**). However, 4MU only inhibited T-cell proliferative responses if treatment began prior to initial antigen presentation (at 0 hours) and not later (at 24 hours)(**Fig. 3J**).

Early activation of antigen specific CD4+ T-cells, as reflected by CD69 expression, was also diminished (**Fig. 3K;** gating shown in **Supplemental Fig. 2C**). T-cell production of the inflammatory cytokines IL-4-, IL-2, and IFNg was likewise decreased, as seen upon intracellular staining of T-cells (**Supplemental Fig. 3**). Importantly, this reduction in activation due to 4MU did not adversely impact the viability of either DCs or T-cells measured 24 hours after addition of 4MU (**Fig. 3L, M**). Together these data suggest that HA inhibition prevents T-cell activation and proliferative responses but only when present at the time of initial antigen presentation.

### 4MU treatment prevents immune synapse formation in vivo

We next sought to understand how interactions between DC and CD4+ T-cell are impacted by 4MU *in vivo*. To this end, we performed 2-photon intravital microscopy and observed the interactions of adoptively transferred ovalbumin (OVA) loaded DC (labeled with celltrace violet) and both OVA-specific, OT-II CD4+ T-cell responders (labelled with SNARF-1) as well as wild type (WT) polyclonal CD4+ T-cells (labelled with Oregon Green). A schematic of this protocol is shown in **Fig. 4A**. Videos (**Supplementary Videos 1**,**2)** taken 16 hours post transfer were used to compare how DC/T-cell interactions were impacted by 4MU. Representative static images from these videos are shown in **Fig. 4B, C**. We observed in mice fed control chow, DCs loaded with OVA preferentially bound to OVA-specific OT-II T-cells, while mice that received 4MU chow OT-II T-cells and polyclonal T-cells bound with equivalent frequency to DC (**Fig. 4D**). Moreover, when we assessed the percentage of labelled DC that were clustered with T-cells, this was significantly decreased on 4MU (**Fig. 4E**). These data indicate that 4MU treatment prevents DC/T-cell interactions *in vivo*.

**Figure 4.**
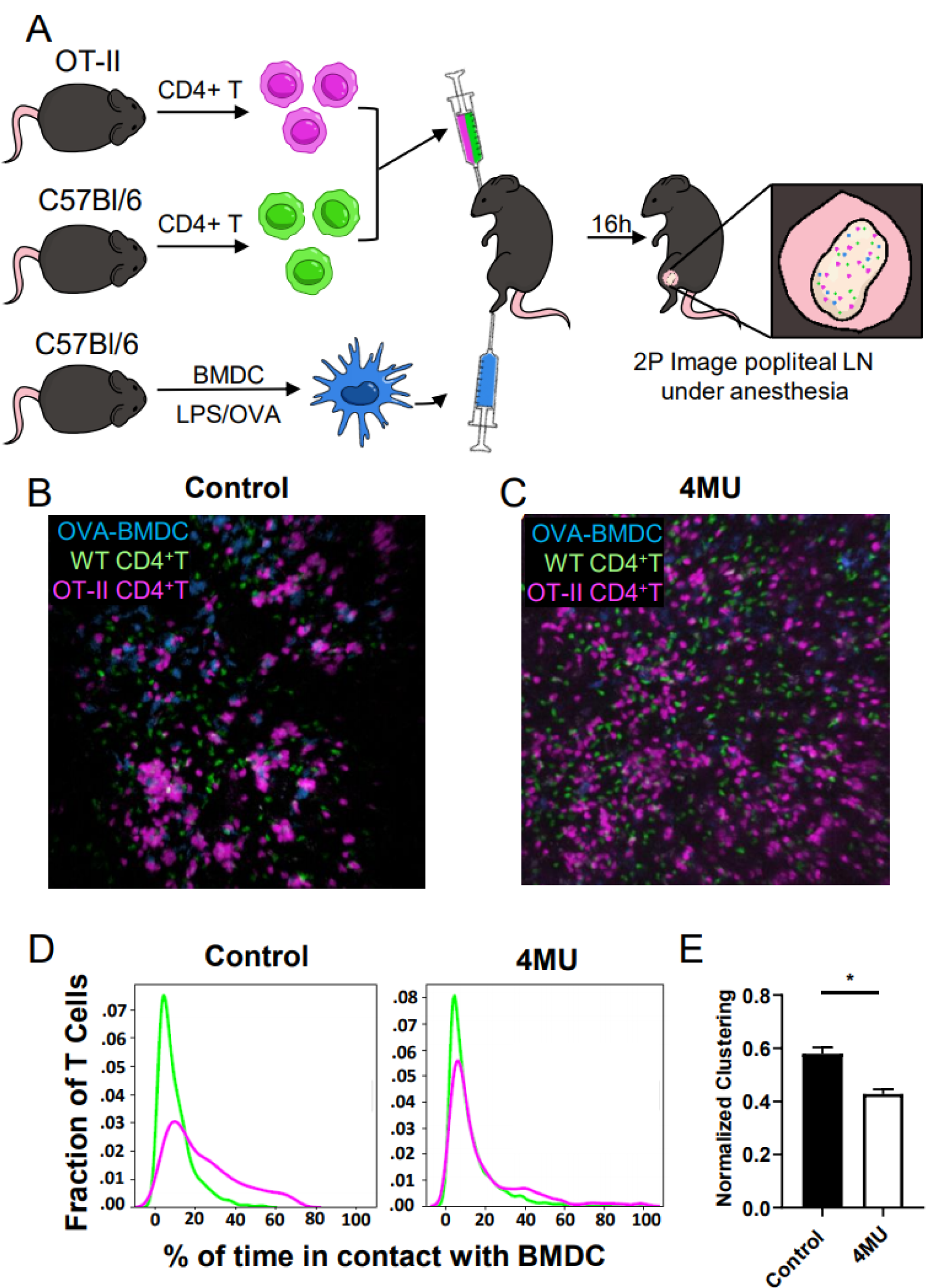
HA contributes to immune synapse formation *in vivo*. A-D. C57Bl/6 animals were pretreated with 4MU or control chow for 14 days prior to an adoptive transfer of labeled CD4+ T-cells and OVA loaded, labeled BMDC. 16 hours after transfer, 2-photon microscopy was used to video the interactions of DC and T-cells within the draining popliteal LN. A. Schematic for experiment. Still images of mice on control (B) or 4MU chow (C) are shown. From video footage (shown in supplemental video 1A and 1B) the percentage of time pink OT-II T-cells spend in contact with BMDC was calculated (D), and is shown in comparison to C57Bl/6 control T-cells in green. (E) Clustering of OT-II T-cells (Pink) to BMDC was determined by normalization to C57Bl/6 T-cells (Green). Data shown are for >500 lymphocytes and >100 DC, and representative of three separate experiments.

### 4MU treatment prevents antigenic responses upon immunization

To investigate how 4MU-induced changes to HA influence *in vivo* T-cell proliferative responses to antigen, we treated C57Bl/6J animals with 4MU in chow or control chow prior to adoptive transfer of OT-II T-cells labeled with a proliferation dye. These animals were then immunized with OVA to stimulate the adoptively transferred cells. Three days later, spleen and draining LN were collected from recipient mice and the proliferation of previously transferred T-cells was assessed via proliferation dye dilution measured by flow cytometry. A schematic of this protocol is shown in **Fig. 5A**. The flow cytometry gating scheme used is shown in **Supplemental Fig. 2D**. We observed a decrease in proliferation of antigen specific T-cells in mice on 4MU compared to controls in both the draining LN and spleen (**Fig. 5B, C**).

**Figure 5.**
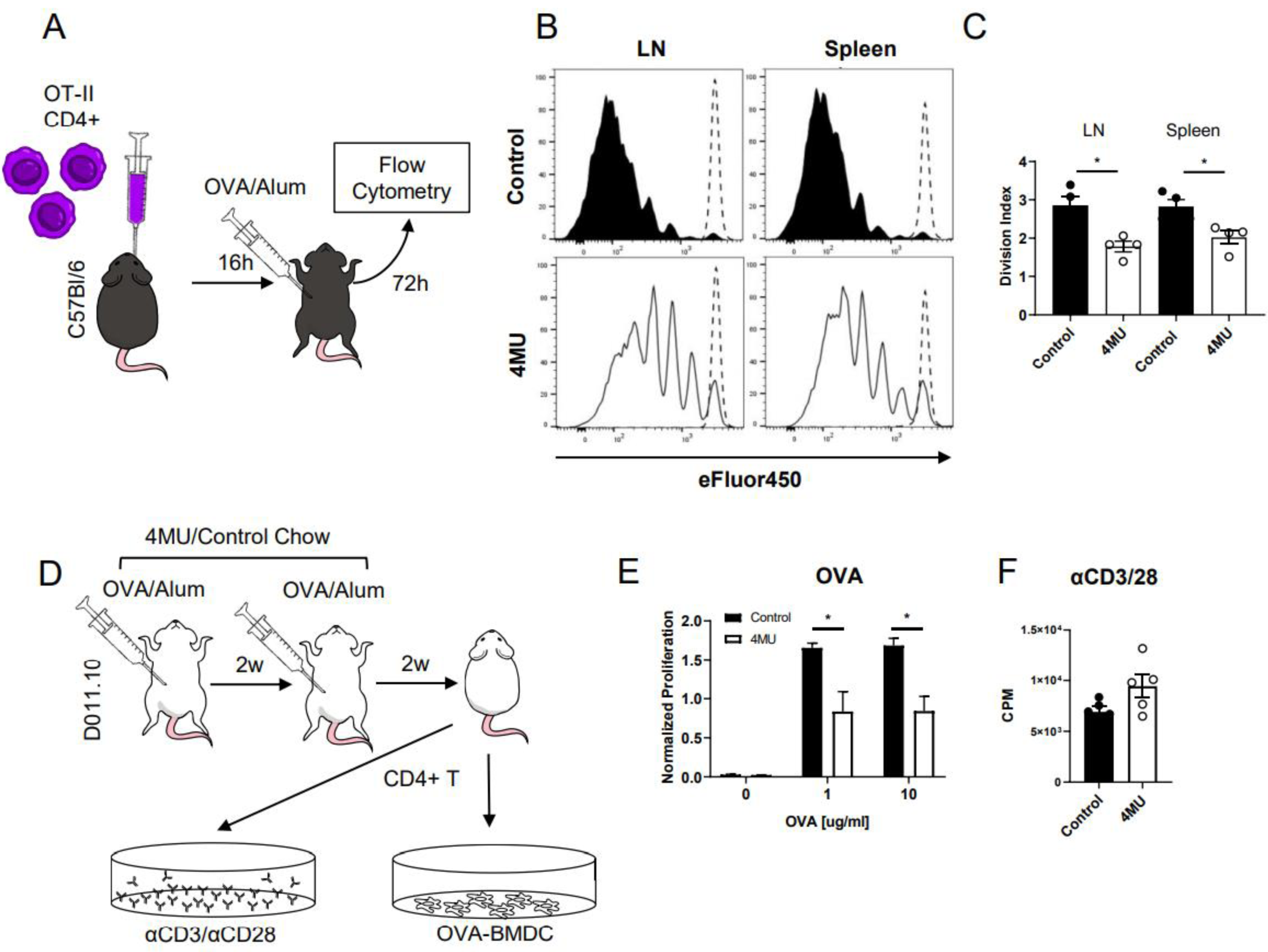
HA contributes to *in vivo* antigen specific responses and development of memory. A-C. 4MU effects on antigen presentation were assessed in an *in vivo* immunization model. A. Schematic for this experiment. 2⨯10 ^6^ ef450 labeled OTII CD4+ T-cells (CD45.1) were transferred into a C57Bl/6 recipient which was then immunized with OVA. After three days, proliferation was assessed via flow cytometry. B. Example of ef450 proliferation of recovered CD45.1 cells. C. Division indices averaged for n=3-4 mice per group. D-F. 4MU effects on recall responses of T-cells isolated from a TCR transgenic mouse (DO11.10) immunized against OVA in the setting of 4MU treatment. Mice were immunized *in vivo* twice to OVA and given time to resolve their inflammation. CD4+ T cells were isolated following sacrifice and the proliferation response to OVA loaded BALB/c BMDC was assessed *ex vivo* after 72 hours via tritiated thymidine uptake. (D) Schematic of the experimental protocol used. E. Proliferation of isolated CD4+ T-cells in response to OVA-peptide pulsed DC, normalized by each animals proliferative response to nonspecific αCD3/αCD28 activation. F. Raw counts for response to αCD3/αCD28. Data collected from n=4-5 animals and measured in triplicate. Representative of three separate experiments. P<.05 as measured by unpaired t-test or one-way ANOVA with multiple comparisons where applicable.

To interrogate the effects of 4MU on the formation of T-cell antigenic memory following immunization, we performed an *ex vivo* re-challenge experiment. We pre-treated OVA-TCR transgenic DO11.10 mice with either 4MU or control chow and then immunized them against OVA twice. These animals were subsequently sacrificed after the primary immune response had resolved, CD4+ T-cells were collected, and these were activated *in vitro*. Activation was done using either freshly generated (never exposed to 4MU) DC pulsed with OVA to provide antigen specific activation, or αCD3/αCD28 to provide antigen independent activation. A schematic of this model is shown in **Fig. 5D**. We observed a strong reduction in the recall response to OVA in cells isolated from animals that were on 4MU during immunization as compared to control animals, while response to αCD3/αCD28 was not impaired (**Fig. 5E,F**). This reduction in antigen specific proliferation is echoed by a reduction in early activation (CD69 expression at 5 hours) (**Supplemental Fig. 4A**). To rule out the possibility of a direct inhibitory effect on memory T cell health by 4MU, we treated mice temporarily with 4MU after the immunizations but before sacrifice and *in vitro* re-challenge. This 4MU intervention group showed no decrease in antigen specific activation as measured by CD69 expression(**Supplemental Fig. 4A**). To further show that antigen specific memory T-cell health was unimpaired by 4MU, we performed a similar experiment observing the proliferative recall response to OVA-loaded BMDC by CD4+ T cells isolated from immunized animals. The proliferative response was unchanged if 4MU was introduced after the immunizations instead of as a pretreatment(**Supplemental Fig. 4B,C**). These findings echo the effects of 4MU treatment timing on *in vitro* responses (**Fig. 3J**). These data suggest that HA inhibition attenuates T-cell activation, but spares pre-established memory.

To control for the possibility that 4MU treatment adversely impacts the homeostasis of T-cells *in vivo*, we transferred CD4+ T-cells from OT-II mice previously immunized with OVA into recipient animals undergoing treatment with either 4MU or control chow (**Supplemental Fig. 4D**). After five days, we saw no change in death or apoptosis of the adoptively transferred cells in either spleen or draining LNs of these animals (**Supplemental Fig. 4E,F)** respectively. As with our *in vitro* viability study (**Fig. 3L-M**) these data suggest 4MU is not intrinsically toxic to T-cells *in vivo*. These data support our previous immune phenotyping of mice on long term 4MU where we likewise did not observe lymphopenia or leukopenia(37).

Together, these data indicate that 4MU pre-treatment prevents *de novo* antigenic responses to immunization while existing memory cell responses are relatively preserved.

### 4MU suppresses allogeneic APC-mediated T-cell proliferation by human cells

We next asked whether 4MU impacts the allogeneic responses by human cells. To examine this, we performed human mixed lymphocyte reactions (MLR)(51), a commonly used allogeneic model of hematopoetic stem cell transplantation. We observed that 4MU prevented allogeneic T cell proliferation in this system (**Fig. 6A**) in a titratable manner. Notably, this effect was seen for six different crosses of different human subjects (**Fig. 6B**), suggesting the inhibitory effect repeats across individual donors and is not dependent on HLA status. Together, these data suggest that HA inhibition reliably dampens human allogeneic T-cell responses.

**Figure 6.**
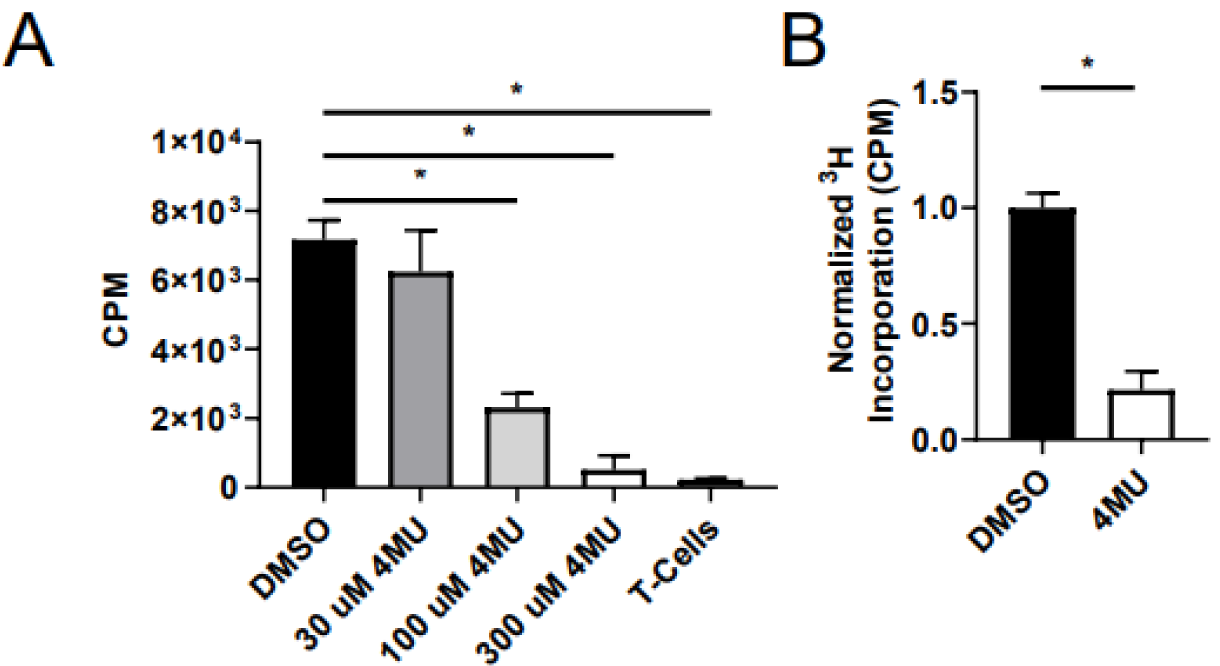
4MU inhibits APC-mediated T-cell proliferation and promotes Treg in a model of graft-versus-host disease. MLR were prepared using monocyte-derived dendritic cells (mDCs) and T-cells isolated from healthy human donors. Cells were cultured for 5 days prior to addition of tritiated thymidine for 24 hours, then measured. (A) A titration of 4MU as well as DMSO control was used. One representative allogeneic cross is shown. (B)Pooled data for 300 µM 4MU condition across 6 different human donor crosses and normalized to control for each cross. Samples were measured in triplicate and p<.05 using unpaired t-test or one-way ANOVA with multiple comparisons where appropriate.

### 4MU treatment delays antigen specific and allogeneic tissue rejection

Given that 4MU impairs antigen recognition by T-cells and promotes Treg, we wondered if 4MU could prevent rejection of allogeneic tissues during transplantation. To examine this, we adoptively transferred OVA-reactive DO11.10 CD4+ T-cells into RIPmOVA recipients (a schematic of this protocol is shown in **Fig. 7A**). As the adoptively transferred cells will attack the OVA-expressing pancreatic islets, the antigen specific response can be tracked by measuring the recipient’s blood glucose. Loss of beta cells will quickly lead to glycemic dysregulation. We observed that pretreatment of the recipient animals with 4MU prevented diabetes for up to 12 weeks after adoptive transfer (**Fig. 7B, C**). However, when 4MU was initiated post-T-cell transfer no protective effect was observed (data not shown).

**Figure 7.**
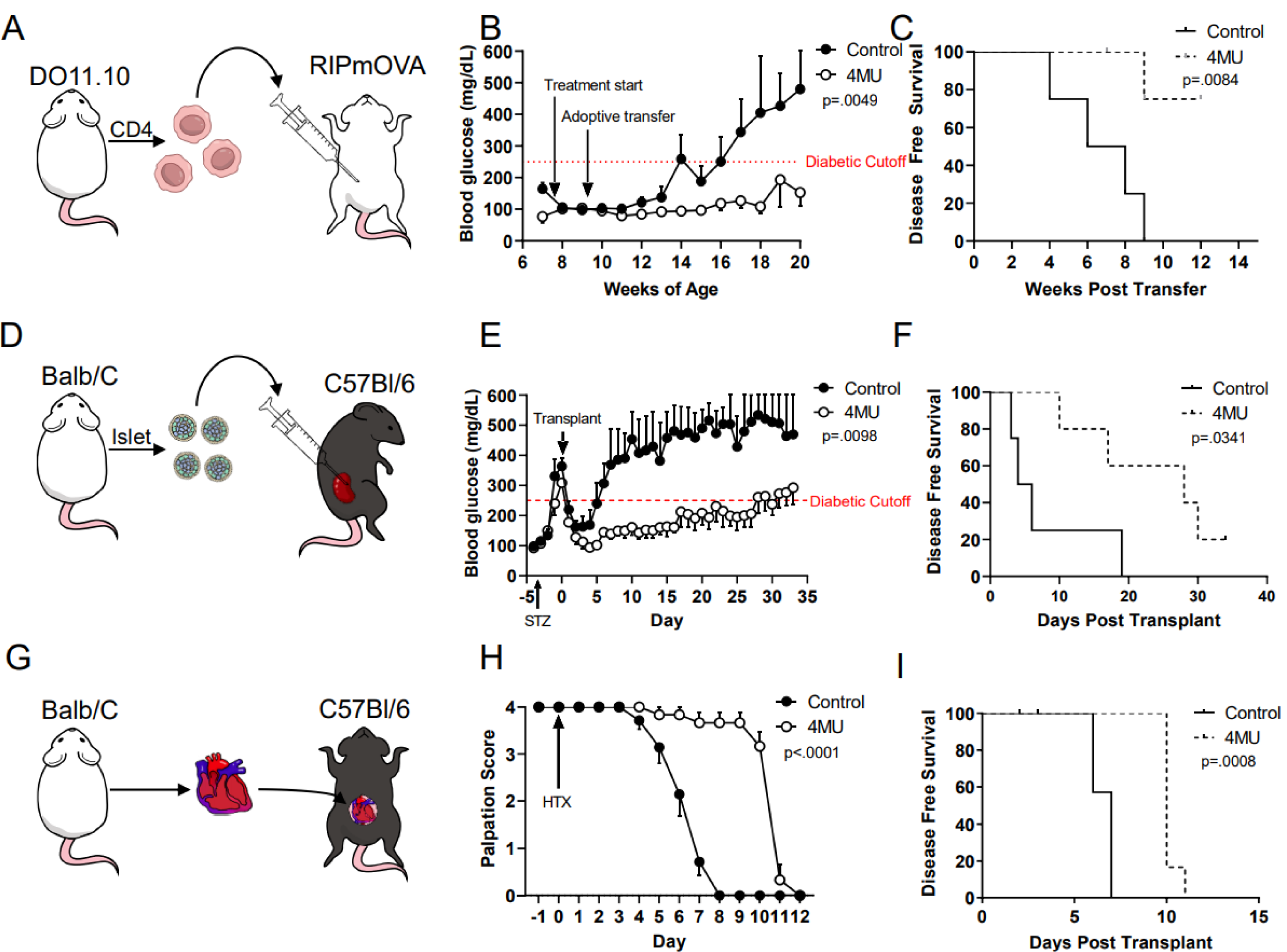
4MU treatment delays rejection of an antigen specific and allogeneic transplantation. (A-C) DO11.10 CD4+ T-cells were adoptively transferred into RIPmOVA recipients, and blood glucose was tracked for 20 weeks to monitor destruction of islets by antigen specific response. (A) A schematic showing the protocol used for the experiments in panels (A-C). (B) Blood glucose levels showing time to diabetes as measured by a blood glucose of 250 mg/dL comparing animals pretreated with 4MU or control chow two weeks prior to adoptive transfer. (C) Kaplan-Meier curve comparing time to diabetes in control and 4MU pretreated animals. (D-F) Adoptive transfer of BALB/C islets into diabetic C57Bl/6 animals, pretreated with 4MU or control chow. (D) Schematic of protocol. In brief, C57Bl/6 transplant recipient animals were treated with STZ to induce diabetes. Four days later, a transplant of BALB/c islets under the kidney capsule of the diabetic animals was done. (E) Daily blood glucose monitoring was used to show the rejection of the MHC mismatched islets. with (F) time to diabetes in the 4MU vs control treated recipients. (G-I) Cardiac allotransplants from BALB/c to C57Bl/6 recipients. (G) A schematic of the protocol used. In brief, mice were treated with control or 4MU chow for 14 days prior to HTX, with engraftment onto abdominal aorta. (H) Mice were then monitored for rejection via palpation of the graft, and detection of contractile activity. (I) Kaplan-Meier curve comparing 4MU or control treated animals. Data are each collected from n= 4-5 animals/group. Blood glucose curves and palpation scores were analyzed by two way ANOVA with multiple comparison tests. Kaplan-Meier curves were compared with Mantel-Cox or Gehan-Breslow-Wilcoxon tests where appropriate.

We then asked whether 4MU could delay rejection in an allogeneic model of pancreatic islet transplantation. C57Bl/6J animals were treated with streptozotocin (STZ) to ablate islets. Three days later, once diabetes was confirmed by blood glucose, islets isolated from MHC mismatched BALB/c animals were adoptively transferred into the kidney capsule of the STZ treated animals (a schematic of this protocol is shown in **Fig. 7D**). We observed that time to diabetes was extended via measurement of blood glucose when mice were pretreated with 4MU chow (**Fig. 7E, F**). Of note, 4MU initiation post-transplant was not assessed because mice require two weeks to reach therapeutic drug levels on 4MU (63) and this was not feasible given the otherwise rapid rejection of allogeneic tissues in this model.

We confirmed this finding using yet another model of transplantation, allogeneic heart transplantation(52, 53). In brief, we pretreated C57Bl/6J recipient animals with 4MU or control chow prior to heterotopic engraftment of an MHC mismatched BALB/c donor heart onto the aorta and inferior vena cava of the recipient (a schematic of this protocol is shown in **Fig. 7G**). Rejection was determined by directly monitoring the heartbeat of the donor heart. We found that heart graft survival was strongly increased in mice that received 4MU chow (**Fig. 7H, I**).

### A model for how HA contributes to antigen presentation and lymphocyte polarization

We conclude that HA plays a critical role in establishing the immune synapse between DC and T-cells and that this is necessary for the development of antigenic responses while 4MU inhibits this. Instead 4MU causes weaker levels of antigenic stimulation, thereby preventing the inflammatory CD4 T cell response that initiates organ rejection. Together, our data suggest that 4MU or other treatments that target pericellular HA may promote allogeneic graft survival in MHC mismatched recipients.

## Discussion

We report that inhibition of HA synthesis with 4MU prevents antigenic responses *in vitro* and *in vivo*. These effects on antigen presentation were observed only when 4MU was added prior to initial antigen presentation but not later, consistent with 4MU-mediated inhibition of *de novo* antigenic responses.

These effects on antigen presentation are consistent with previous reports that 4MU treatment prevents disease progression in several models of autoimmunity(37)(55)(56)(38), as autoimmune diseases reflect dysregulated adaptive immunity. Further, 4MU was also previously reported to induce Foxp3+ Treg; this too is consistent with 4MU effects on antigen presentation, as weaker and shorter interactions promote the induction of Treg(57, 58, 59). These data do not rule out roles for HA in signaling or other metabolic or cellular processes previously attributed to 4MU but instead suggest that these occur together with effects on DC/T-cell interactions.

Our data are consistent with a role for HA in stabilizing interactions between DC and T-cells. Within lymphatic tissues, HA is abundant within the germinal centers and interfollicular zones where antigen is presented and associated with cell types involved in antigen presentation, namely DCs, macrophages, monocytes, and a subset of B cells. Moreover, treatment with 4MU resulted in an interruption of DC/T-cell interactions both *in vitro* and *in vivo*. These data echo similar results previously reported for PEP-1 and hyaluronidase treatment(28, 54). However, while 4MU can be administered orally, agents like hyaluronidase require systemic IV infusion and would be impractical for chronic administration therapeutically.

Building on these findings, we find that 4MU treatment delays rejection of transplanted allogeneic pancreatic islets in mice and suppresses allogeneic T-cell activation in human mixed lymphocyte reactions. In light of these findings, it might be possible to treat patients prior to and following transplantation, perhaps as an adjunct to conventional immunosuppressants. While tremendous progress has been made over the preceding decades in developing safe and effective immunosuppressive agents for solid organ transplantation(61), there remains a great need for novel treatment strategies and new therapeutic targets(62). 4MU is already an approved drug with a benign safety profile(36, 37, 57, 63). Incorporating this agent into existing regimens may allow proportionately less of other, potentially more toxic therapeutic options to be used.

These studies raise several questions that remain to be addressed. The optimal dosages and treatment regimens for HA synthesis inhibition in humans are unknown and the potential interactions between 4MU and existing immunosuppressants are unclear. Our current efforts are directed at obtaining a detailed understanding of HA’s contribution to immune synapse formation and the role of costimulation through the primary HA receptor CD44 in these responses. However, as our focus was on 4MU in this study, those subjects were beyond the scope of these current efforts.

In conclusion, we propose that pericellular HA is a novel frontier in both our understanding of antigenic priming and that it may be possible to target these structures using 4MU for therapeutic benefit in transplant and other settings.

## Materials and Methods

### Mouse Handling

Foxp3-GFP C57Bl/6 mice were the kind gift of Dr. Alexander Rudensky. OTII and DO11.10 mice were purchased from The Jackson Laboratory (Bar Harbor, Me). OTII mice were crossed to Foxp3-GFP mice as well as to the C57Bl/6 CD45.1 congenic strain from The Jackson Laboratory (Bar Harbor, Me). RIPmOVA mice were available at the Benaroya Research Institute (Seattle, WA) and crossed to DO11.10 mice purchased from The Jackson Laboratory to create the DORmO mice. C57Bl/6J and BALB/c mice were purchased from The Jackson Laboratory. All mice were maintained in specific pathogen-free AAALAC-accredited animal facilities at the Benaroya Research Institute and Stanford University and handled in accordance with institutional guidelines.

### Experimental Details

#### Human blood samples

Human peripheral blood samples were obtained from healthy volunteers with informed consent, participating in a research protocol approved by the institutional review board of Stanford University. Human PBMC were isolated as previously described(64). CD4+ T-cells were isolated using a RosetteSep(tm) kit (Stemcell Technologies, Vancouver, BC) according to manufacturer’s instructions. Cells were cultured in RPMI (Gibco, Waltham, Massachusetts) supplemented with 10% Fetal Bovine Serum (Hyclone, Logan, Utah), 100 μg/ml Penicillin, 100 U/ml Streptomycin, and 1 mM sodium pyruvate (Invitrogen, Carlsbad, CA) as previously described(64). Human monocyte-derived DC were generated as previously described(65).

#### Isolation and culture of mouse leukocyte populations

Total leukocytes were isolated from LN and spleen from 8 to 12 week old mice as previously described(64). CD4+ T-cells were isolated from the pooled cell suspensions using an EasySep(tm) Mouse CD4+ T Cell Isolation Kit from Stemcell Technologies (Vancouver, Canada) following the manufacturer’s instructions. Foxp3+ Treg induction protocols were as previously described(43). Polyclonal T-cell activation was performed as previously described(66), pre-coating plates with 10 µg/ml αCD3 (145-2C11, Biolegend) in PBS overnight at 4°C. Coated plates were washed and cells were plated with 1 µg/ml soluble αCD28 antibody (37.51, Biolegend). Bone marrow-derived dendritic cells (BMDC) were generated as previously described(65). DCs were activated overnight with 1 ug/ml LPS (Sigma) along with OVA peptide 323-339 (Invivogen) for antigen specific activation. For polyclonal activation via APC, BMDC Fc receptors were pre-loaded with 2.5 ug/ml αCD23 (Biolegend) and .5 ug/ml αCD28 (Biolegend) for two hours before culturing with target CD4+ T-cells. Antigen processing assays were performed according to manufacturer’s protocol for DQ-OVA (Thermo Fisher Scientific).

### Flow Cytometry

Cells were stained and flow cytometry was performed as previously described(64). Analysis for this project was done on LSR II (BD Biosciences) instruments in the Stanford Shared FACS Facility, as well as a Cytek NL-3000. Analysis, including generation of tSNE plots, was done on Flowjo v10.

Human flow cytometry experiments used the following fluorochrome-labeled antibodies: CD14 (HCD14), CD20 (2H7), CD19 (H1B19), CD56 (HCD56), CD25 (BC96)(all Biolegend), and CD4 (S3.5), CD45RA (MEM-56), CD45RO (UCHL1)(all Life Technologies, Carlsbad, CA), and CD44 (G44-26, BD Biosciences, San Jose, CA).

Mouse flow cytometry experiments used the following fluorochrome-labeled antibodies: TCR Vβ (H57-597), CD19 (6D5), I-A/I-E (M5/114.15.2), F4/80 (BM8), CD11b (M1/70), CD11c (N418), CD14 (Sa14-2,), CD4 (GK1.5), CD69 (H1.2F3), CD45.1 (A20), CD45.2 (104), Vα2 (B20.1), CD44 (IM7,), CD62L (MEL-14)(all Biolegend) and CD49b (DX5), CD8α (53-6.7), Foxp3 (FJK-16s)(all ThermoFisher) and CD3ε (500A2, BD Biosciences).

To detect surface HA, cells were stained overnight in 50 µg/ml biotinylated HABP (EMD Millipore) in T-cell media. Cells were washed and stained with a secondary Alexa 647 biotin (ThermoFisher Scientific) at 1 µg/ml alongside traditional surface antibodies at 4°C for 30 minutes, prior to fixation in 10% NBF for 10 minutes at room temperature before washing.

For cell proliferation studies, Cell Proliferation eFluor 450 (ThermoFisher Scientific) was used according the manufacturer’s protocol.

### 4MU treatment

In culture, 4MU was dissolved in DMSO and was added at 100 µg/ml (approximately 300 µM). For *in vivo* studies, 4MU (Alfa Aesar, Haverhill, MA) was pressed into the mouse chow by TestDiet® as previously described(63). Mice were initiated on the 4MU chow at 6 weeks of age and were maintained on this diet until euthanized, unless otherwise noted.

### Scanning Electron Microscopy (SEM)

BMDC were isolated as described above, and grown on fibronectin-coated glass slides for 24 hours, were fixed overnight in 4% paraformaldehyde with 2% glutaraldehyde in 0.1 M sodium cacodylate buffer (pH 7.4). The slides were gently washed twice with the same buffer, and then post-fixed in 1% aqueous osmium tetroxide (OsO4) for one hour. Samples were then washed twice in purified water and dehydrated in an increasing ethanol series (50%, 70%, 80%, 90%, 100%, 100%, 5 min each). Finally, the specimens were critical-point dried (CPD) in liquid CO _2_, in a Tousimis 815B critical-point dryer (Tousimis, MD). CPD-dried samples were mounted on SEM stubs with adhesive copper tape and sputter-coated with 4 nm of Au/Pd film (4 nm in thickness) in a Denton Desk II machine (Denton Vacuum, NJ). The prepared SEM samples were examined with a Zeiss Sigma field emission SEM (FESEM) (Carl Zeiss Microscopy, NY) operated at 2-3 kV, using InLens SE detection.

### T-cell - DC *In Vitro* Binding

*In vitro* DC/T-cell binding assays were performed as previously described(28). In brief, BMDC were stained using SNARF-1 (Invitrogen) according to manufacturer’s protocol. After this 1⨯10 ^5^ BMDC per condition were cultured on a collagen (Fisher Scientific) coated6-well plate (Corning, Corning, NY) overnight. BMDC were maintained under these culture conditions overnight alongside 1 µg/ml LPS (Sigma-Aldrich) and 1 µg/ml αCD3. The plates were gently washed two times to remove any conditioned media and dead APC. Naïve CD4+ T-cells were then labeled with 50 μM, carboxyfluorescein succinimidyl ester (CFSE)(Invitrogen). 1⨯10 ^5^ CFSE labeled CD4+ T-cells were added to the plates. After incubation for 4 hours the plates were washed in PBS supplemented with 1mM CaCl2 amd 1 mM MgCl2, followed by fixation with 10% NBF for 30 minutes, washed a second time in PBS, and stored in PBS until analysis. Analysis of binding was performed as follows. For each condition, at least 10 non-overlapping fields were photographed using a digital camera (Diagnostic Instruments, Sterling Height, MI) attached to a Leica DM-IRB microscope (Leica Microsystems, Wetzlar, Germany). Spot Software 4.5 (Diagnostic Instruments; Sterling Heights, MI) was used for analysis. For each field photographs were taken using the excitation laser at 488 nm and at 568 nm in order to capture binding of both CFSE-labeled T-cells as well as BMDC labeled with SNARF-1. For each field the two images were then merged using software in order to provide an assessment of clustering involving both T-cells and DC.

### Immunocytochemistry, Quantification of HA and the HA synthase isoforms

HA immunocytochemistry was performed as previously described(31). Periucellular HA quantification was performed using a radiolabeled glucosamine incorporation assay, as previously described(28). HAS1-3 quantitative PCR was performed as previously described(37).

### MLR Mixed lymphocyte reactions (MLRs)

Allogeneic monocyte-derived dendritic cells (moDCs) and T-cells were isolated from healthy human donors. moDCs were obtained by isolating monocytes from PBMCs using CD14 MicroBeads (Miltenyi Biotec, Bergisch Gladbach, Germany), following Miltenyi Biotec’s protocol for generation of moDCs. Briefly, monocytes were cultured with 250 U/ml IL-4 (R&D Systems, Minneapolis, MN) and 800 U/mL GM-CSF (R&D Systems) for 6 days. T-cells were obtained from PBMCs using the EasySep Human T-cell Isolation Kit (Stemcell Technologies). MLRs were set up in 96 well plates. Each MLR well contained 5×10 ^4^ moDCs and 2.0⨯10 ^5^ T-cells for a ratio of 1:4 moDCs:T-cells. Prior to plating, moDCs were irradiated at 40 Gy. Positive proliferation control wells were set up by plating 1 Dynabeads Human T-Activator CD3/CD28 (ThermoFisher Scientific) for every 200 T-cells. Negative proliferation control wells were set up by plating T-cells only. Drugs were dissolved in DMSO and added at the specific concentrations (30, 100, or 300 µM) in triplicate. The wells were cultured for 5 days, to which tritiated thymidine was added and proliferation assessed 24 hours later.

### Heterotopic Allogeneic Cardiac Transplants

Heterotopic cardiac transplantations were performed as previous reported(52, 53). Briefly, a BALB/c donor heart was transplanted into recipient’s abdomen by anastomosing the donor aorta to C57Bl/6J recipient’s abdominal aorta and donor pulmonary artery to recipient’s inferior vena cava (IVC) at an end-to-side manner. Cardiac graft function was assessed through abdominal palpation.

### Islet Transplantation

BALB/c mice of 12–24 weeks age were anesthetized, the descending aorta of each anesthetized mouse was transected, the bile duct clamped at its distal (intestinal) end, and a 30-gauge needle was used to inflate each pancreas through the common bile duct with 4 mL of media supplemented with 0.8 mg/mL of collagenase P (Roche, cat. number 11-249-002-001) and filtered at 0.22 μm prior to injection. After all pancreata were processed, the isolated islets were hand-picked, cultured overnight, and picked again the next day before transplant. C57Bl/6 transplant recipient animals received 4MU or control chow for two weeks before being treated with STZ to induce diabetes. To this end, mice were treated with a high dose (200 mg/kg) of STZ made as a stock solution of 7.5 mg/ml STZ in 100 mM citrate, pH 4.2 (prepared and filtered at 0.22 μm immediately prior to intraperitoneal injection). Four days later, a transplant of Balb/c islets under the kidney capsule of the diabetic animals was done. Daily blood glucose monitoring was used to show the rejection of the MHC mismatched islets. with time to diabetes in the 4MU vs control treated recipients.

### Two Photon Intravital Microscopy and Analysis

CD4+ cells were isolated from spleens and LN from C57Bl/6 or OT-II via negative selection as mentioned above. Bone Marrow derived Dendritic cells (BMDC) where cultured according to standard protocol. T-cells and BMDCs were labeled with fluorescent dye (SNARF-1/Oregon Green/Celltrace Violet ThermoFisher) according to the manufacturer’s protocol. We transferred 5×10 ^6^ labeled CD4+ cells T-cells into recipient mice 16 h before imaging by i.v injection. BMDCs where injected intradermally into a footpad 16 h before imaging. Mice were then anesthetized and images were acquired as previously described(38).

Image analysis was performed with Imaris software (Bitplane Inc., Belfast, United Kingdom). 3D velocity of T-cells was calculated from the coordinates of their centroids tracked for the full length of a movie (15–30 min). Time in contact was calculated based on these movies. Clustering was calculated by normalizing BMDC interaction with C57Bl/6 T-cells as a control for clustering interactions. Based on these interactions clustering between BMDC and OTII was calculated.

### Statistical Analysis

Where n is not stated, graphs show a representative experiment of n≥2 assays, with n≥3 technical or biological replicates. The number of mice needed for the *in vivo* experiment was determined using power calculations. All statistical analyses, linear and nonlinear regression analyses were performed using GraphPad Prism (GraphPad Software, Inc. La Jolla, CA). All unpaired Student’s t-tests, ANOVA, Mann-Whitney tests, and Fisher’s exact test were two-tailed. For ANOVA, Holm-Sidakk and Bonferroni multiple comparisons were performed where appropriate and as indicated. Depicted are means with SEM of the replicates unless otherwise stated. Statistical significance was considered p<0.05.

## Supporting information

Graphical Abstract and Highlights

## Supplementary Materials

**Supplemental Figure 1:**
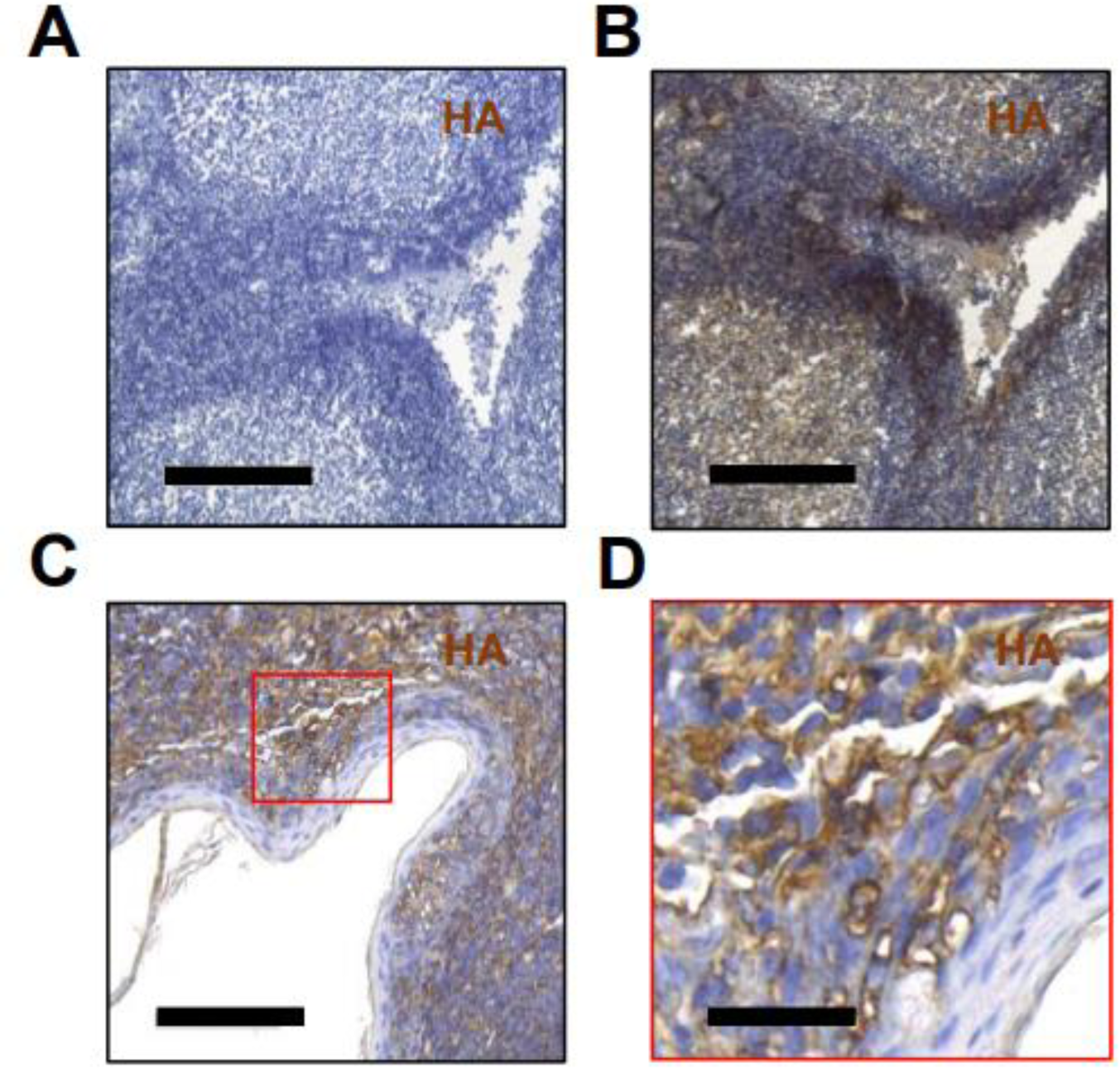
HA is abundant in human lymphatic tissue. Human tonsils were obtained from de-identified, discarded surgical specimens. Representative staining of sections from a single individual is shown for (A) an unstained section, and (B) the same section stained for HABP, (C) An area of dense lymphocytic predominance, and (D) the same area as in (C), now at high magnification. Black bars = 100 µm in (A-C) and 25 uM in (D). Data are representative of tonsil tissue from 3 subjects.

**Supplemental Figure 2:**
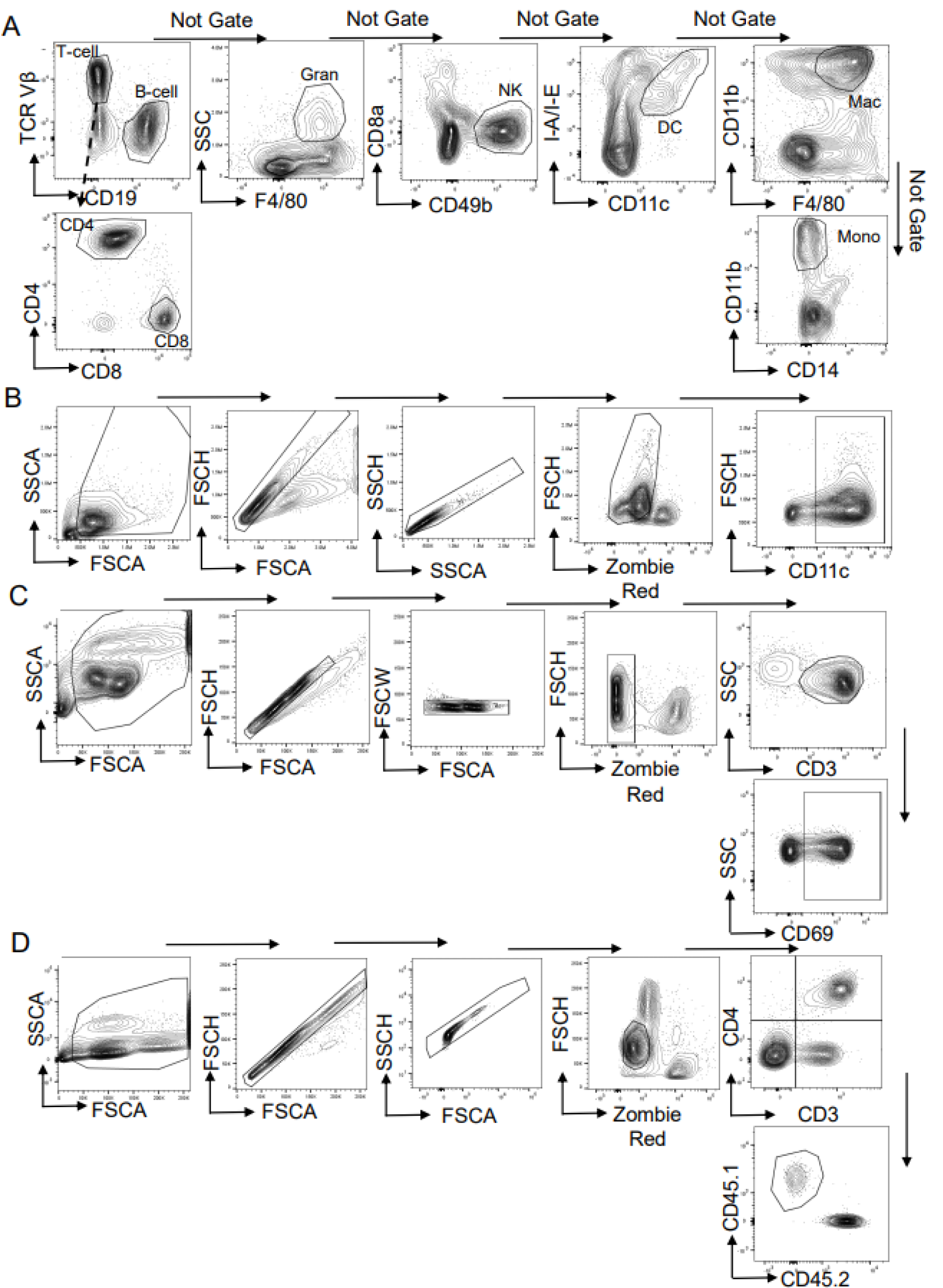
Gating scheme for mouse experiments described in Figures 1-5. A. Gating scheme for figure 1(G) of DORmO secondary lymphatic tissue. Following singlet and live/dead gating, cells were gated by primary lineage markers for relevant cell groups. Gating scheme identifies cell types by common surface markers, then “Not gates” to prevent duplicated cell definitions in other cell types. Data is representative of three separate animals. B. Gating scheme for Figure 2(D-G). Gating scheme for BMDC phenotyping under 4MU treatment. For observing surface phenotypic markers, BMDC were only defined as CD11c+, Zombie Red (viability) negative. C. Gating scheme for Figure 3K. Gating scheme for early activation of CD4+ OTII T-cells by OVA-pulsed BMDC. %Activated CD4+ T-cells after 5 hour incubation was defined as %CD69+ cells of live CD3+ T-cells. D. Gating scheme for Figure 5(A-C). Gating scheme for adoptive transfer of congenic (CD45.1), efluor450 labeled CD4+ T-cells from an OTII animal. To observe division of adoptively transferred cells, gating was done on live, CD3+, CD4+, CD45.1, CD45.2-singlets. Fit curves created on Flowjo v10 were used to determine division index of these cells per recipient.

**Supplemental Figure 3:**
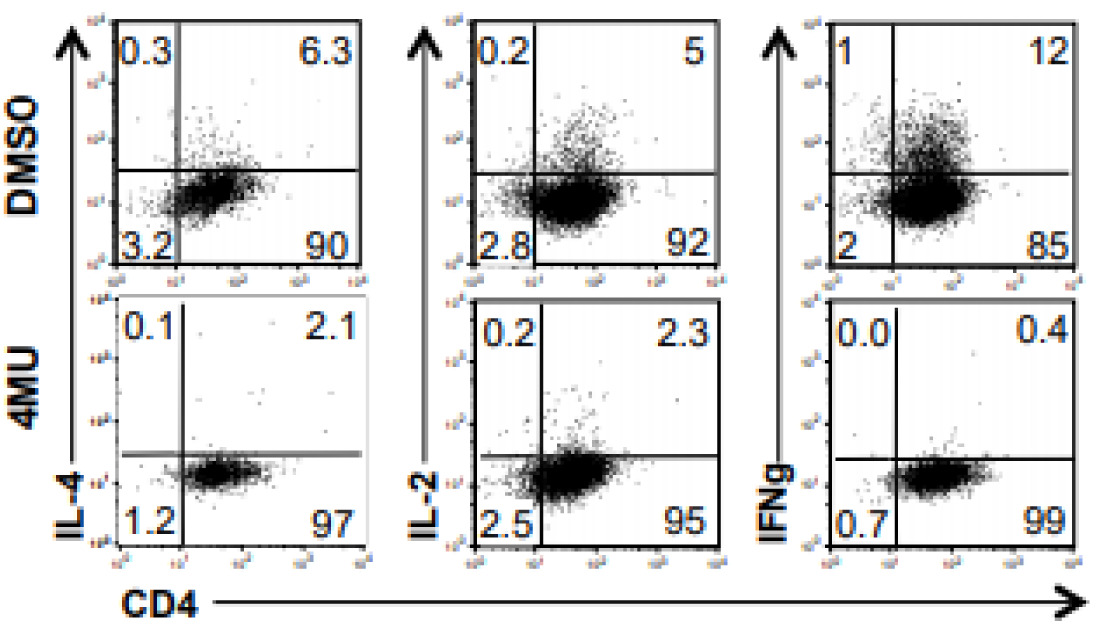
Intracellular cytokine staining during 4MU treatment. Human mDC were used to stimulate CD4+ T-cells combined with αCD3/α28 antibodies. After 72 hours, cells were stained for intracellular cytokines. Data are representative of 3 independent experiments performed using cells from different donors.

**Supplemental Figure 4:**
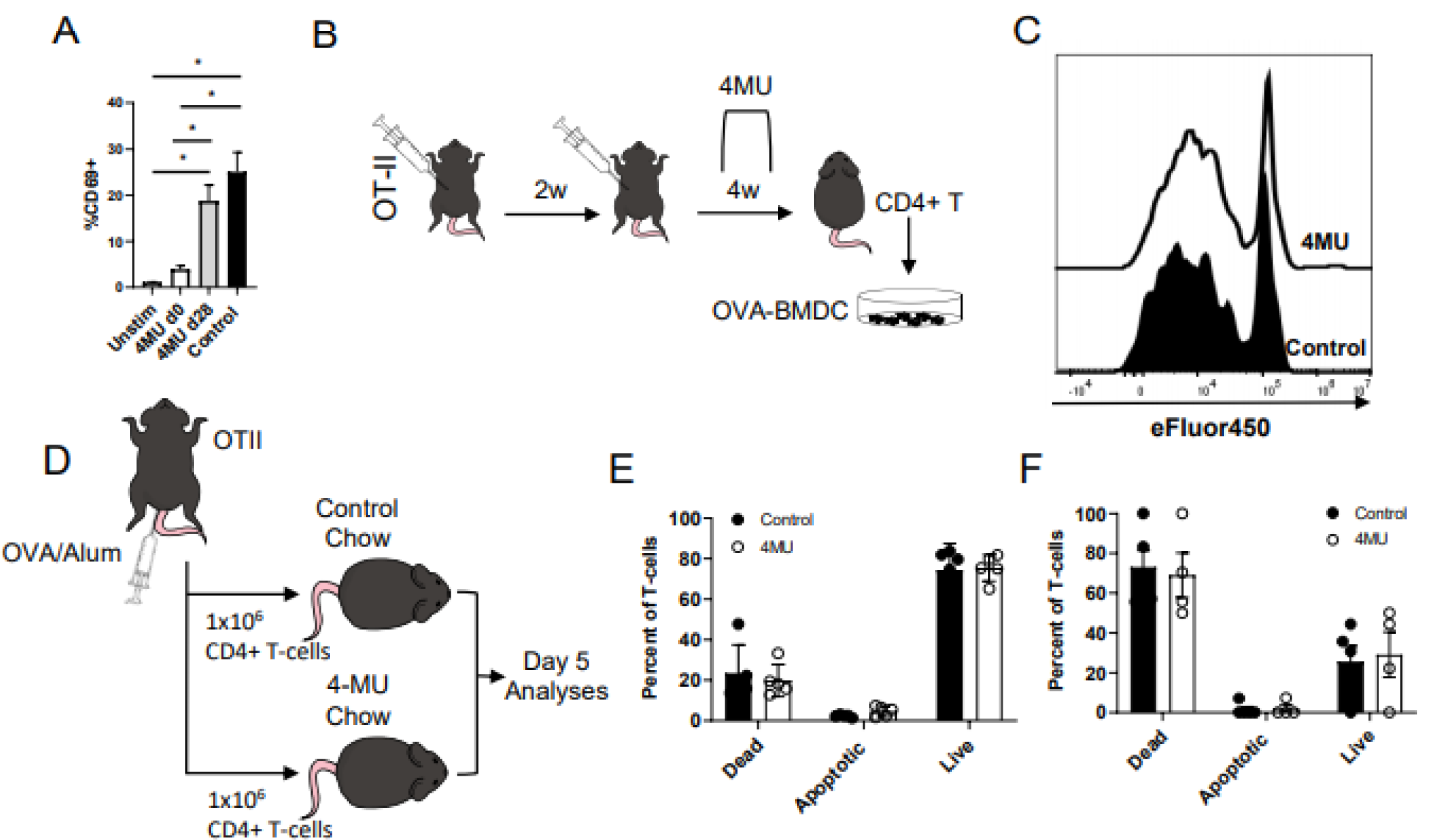
4MU does not impact the viability of previously activated T-cells. A. Cells were isolated from OVA-immunized OT—II animals in the same manner and groups as Figure 5D-F. Animals were allowed two weeks following second immunization to resolve inflammation before sacrifice and isolation of CD4+ T-cells. 5 hours after *ex vivo* stimulation by OVA-loaded C57Bl/6 DCs, T-cells were evaluated for CD69 expression via flow cytometry. B,C OT-II animals were immunized to OVA and isolated CD4+ T cells were stimulated *ex vivo* via OVA loaded BMDCs for three days. Proliferative response was measured via dilution of the proliferation dye ef450. Histogram is representative of results from four separate animals measured in triplicate D-F 4MU does not impair T-cell homeostasis following *in vivo* adoptive transfer of pre-activated T-cells. D. Experimental scheme. CD45.1 OTII mice were immunized with Ovalbumin and CD4+ T-cells were isolated from spleen and draining LNs of immunized mice. One million cells were transferred to each recipient C57Bl/6 CD45.2 mouse, retro-orbitally. Recipient mice were either on control chow or on 4MU chow for 14 days prior to transfer. 5 days post transfer spleens and LNs were harvested from recipient mice. B,C Viability of donor T-cells represented as % of live, apoptotic or dead is quantified for spleen (E) and peripheral LN (F) of recipients. Error bars represent standard error of the mean. N=4 per group. No differences are significant by an unpaired t-test.

**Video S1 Representative video of control mouse, 2 photon imaging, post-processing**. After videos are collected, images are segmented into planes on Imaris. Interacting objects are shown frame by frame with OT-II CD4+ T-cells in pink, polyclonal CD4+ T-cells (C57Bl/6) in green, and OVA-loaded BMDC in blue. Mice were injected retro-orbitally with OT-II T-cells and polyclonal T-cells the afternoon before imaging. LPS and OVA pulsed BMDC were injected into the rear foot pad of the animal at the same time. Starting at t=16 hours, videos ranging from 15-30 minutes were collected.

**Video S2 Representative video of 4MU treated mouse, 2 photon imaging, post-processing**. After videos are collected, images are segmented into planes on Imaris. Interacting objects are shown frame by frame with OT-II CD4+ T-cells in pink, polyclonal CD4+ T-cells (C57Bl/6) in green, and OVA-loaded BMDC in blue. Mice were injected retro-orbitally with OT-II T-cells and polyclonal T-cells the afternoon before imaging. LPS and OVA pulsed BMDC were injected into the rear foot pad of the animal at the same time. Starting at t=16 hours, videos ranging from 15-30 minutes were collected.

## Acknowledgments

This work was supported by grants from the NIH (R01 DK096087-07, R01 DK114174-01A1, and U01 AI101984 to PLB. This work was also supported by Juvenile Diabetes Research Foundation (JDRF) grant 3-PDF-2014-224-A-N (to N. N.) and the Stanford Diabetes Research Center grant P30DK116074, National Institutes of Health, SPARK Translational Research Program Pilot Grant, DFG grants to N.N. (NA 965/2-1) and G.K. (KA 3441/1-1) as well as a Stanford University Bio-X Bowes Fellowship to P.L.M.. We thank the Stanford Shared FACS Facility for technical assistance. The authors have no competing financial interests. N.N., P.L.M., and P.L.B. are listed as inventors of the patents (PCT/US2019/019310) filed by the Board of Trustees of the Leland Stanford Junior University.

