## Supplementary material for "Hyaluronan Synthesis Inhibition Impairs Antigen Presentation and Delays Transplantation Rejection": Graphical Abstract and Highlights

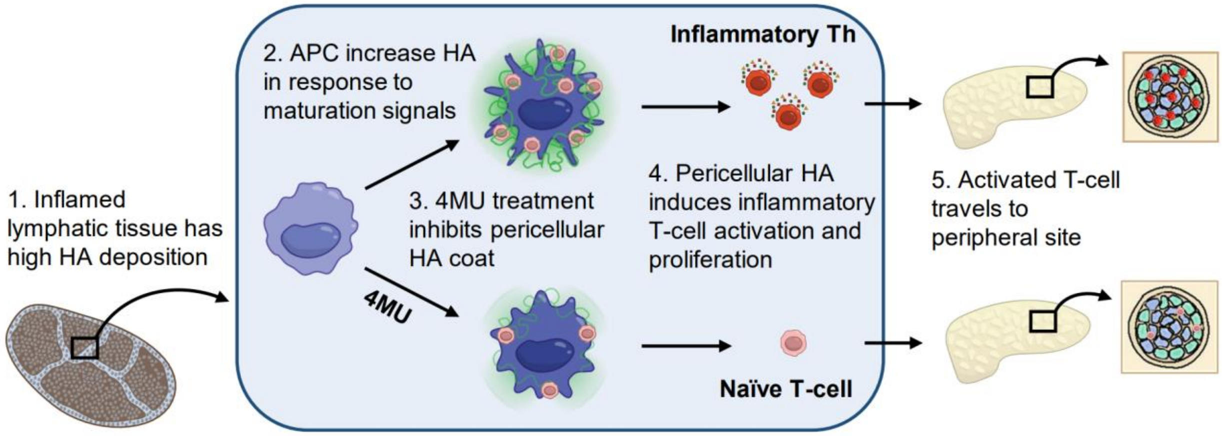


**Highlights**

- Hyaluronan is deposited in secondary lymphoid tissue at sites of inflammation by antigen presenting cells.
- Inhibition of hyaluronan during antigen presentation prevents activation and proliferation of inflammatory CD4+ T-cells.
- Treatment of mice with the hyaluronan inhibitor 4-methylumbelliferone prevents antigen specific T-cell responses and memory response during re-challenge.
- 4-methylumbelliferone inhibits allogeneic T-cell responses and prolongs time to rejection of solid organ transplantation.
